# LEVERAGING THE POWER OF 3D BRAIN-WIDE IMAGING AND MAPPING TOOLS FOR BRAIN INJURY RESEARCH IN MURINE MODELS

**DOI:** 10.1101/2023.04.27.537761

**Authors:** Mehwish Anwer, Jeffrey LeDue, Zefang Wang, Sarah Wang, Wai Hang Cheng, Mariia Burdyniuk, Honor Cheung, Jianjia Fan, Carlos Barron, Peter A Cripton, Mark S Cembrowski, Fabio Rossi, Timothy H Murphy, Cheryl L Wellington

## Abstract

Despite the fundamental importance of understanding impaired brain activity exhibited in post-traumatic epilepsy and other neurological impairments associated with traumatic brain injury (TBI), knowledge of how brain injury affects neuronal activity remains remarkably incomplete. We describe a whole-brain imaging and analysis approach to identify alterations in neuronal activity after TBI as a complementary method to conventional two-dimensional (2D) histological approaches. Here we report an easy-to-follow experimental pipeline to quantify changes in the whole mouse brain using tissue clearing, light sheet microscopy (LSM) and an optimised open-access atlas registration workflow. We validated the outcome of the pipeline using high throughput image analysis software and a secondary atlas registration method. Using the CHIMERA (Closed-Head Impact Model of Engineered Rotational Acceleration) TBI model, TRAP2 mice were subjected to repeated mild TBI or sham treatment followed by tamoxifen injection to lock c-Fos activity after TBI. Brains were SHIELD fixed and passively cleared for imaging of c-Fos+ cells throughout the rostro-caudal axis of the brain using a light sheet microscope equipped with a specialized whole-brain imaging chamber. Volumetric images were stitched and 3D rendered using Arivis Vision4D image analysis software. For quantitative analysis, 2D image stacks were exported to segment c-Fos+ cells and register them to the Allen Mouse Brain Atlas using the BrainQuant3D python package. As a result, c-Fos+ cell counts were estimated throughout the brain and heatmaps were generated. We identified a brain-wide reduction in c-Fos cell density in the TBI group compared to sham controls, indicative of TBI-induced changes in whole brain neuronal activity. Further studies using multi-dimensional imaging coupled with analysis tools will deepen our understanding of post-TBI brain-wide dynamics.

## INTRODUCTION

Traumatic brain injury (TBI) is a leading cause of death and disability with a global incidence of ∼70M (Dewan et al., 2018) that is associated with a high incidence of neurological impairment including post-traumatic epilepsy. TBI induces a myriad of pathological processes resulting in brain-wide damage to cellular activity, axonal connectivity, and vasculature integrity (V. E. Johnson, Stewart, & Smith, 2013; Shlosberg, Benifla, Kaufer, & Friedman, 2010; Smith, Johnson, & Stewart, 2013). Despite the fundamental importance of understanding these processes, our knowledge to date remains remarkably incomplete. In this study, we leverage whole-brain imaging and analysis approaches to quantify altered neuronal activity in mice after TBI using the Closed Head Impact Model of Engineered Rotational Acceleration (CHIMERA) model, which uses biomechanical impacts to reliably induce unrestricted head acceleration and cause diffuse axonal injury, diffuse microvascular injury, inflammation, and blood biomarker and neuroimaging changes similar to human TBI (Bashir et al., 2020; Cheng et al., 2018, 2019; Martens, Cheng, Namjoshi, Cripton, & Wellington, 2016; McNamara, Grillakis, Tucker, & McCabe, 2020; Namjoshi et al., 2014, 2017; W.H. et al., 2019).

Brain tissues collected from animal TBI models fill a critical gap posed by the lack of clinical specimens. The diffuse nature of many TBI pathologies poses considerable challenges to traditional histopathological analyses, as 2D sections could easily miss key affected areas even when a systematic approach to scan the brain using multiple sections per animal is used. To address this needle-in-the-haystack problem, we leveraged light sheet microscopy (LSM) to provide an unbiased analysis of whole brain spatial pathological changes after CHIMERA TBI. LSM is an efficient way to study the diffusely injured brain as, compared to routine 2D histological methods (Ahrens, Orger, Robson, Li, & Keller, 2013; Hillman, Voleti, Li, & Yu, 2019), LSM preserves native tissue morphology thereby enabling mapping of cortical and subcortical damage after TBI. To accelerate unbiased multi-dimensional assessment of TBI-induced neuronal changes, we optimized a tissue clearing and LSM protocol to assess altered neuronal activity. We carried out TBI in Fos^2A-iCreER^ TRAP2 mice (DeNardo et al., 2019), that express a trappable red fluorescent protein variant (TdTomato) under the control of the immediate early gene (IEG) *Fos* promoter. The c-Fos expression at stimulus epochs is TRAPed within few hours of tamoxifen injection and as the reporter protein expression with TRAP2 is permanent, analysis of TRAPed cells can be performed long after TRAPing has occurred, thereby allowing us to visualize activated neurons after TBI.

Advances in tissue clearing, immunolabelling, and volumetric fluorescent imaging provide the methodology to extract brain-wide imaging datasets of IEG expression. However, there is a need for computational tools that streamline their analysis and include human checkpoints for orthogonal cross-validation of automated approaches (Newmaster, Kronman, Wu, & Kim, 2022). We thus optimized BrainQuant3D, an open-source python-based package (Schneider et al., 2019), which builds on existing registration and automated cell counting methods (Renier et al., 2016), to facilitate whole-brain c-Fos mapping in mice and optimize a workflow that uses whole brain imaging tools and downstream data processing methods to enable brain-wide analysis of target cellular populations in the murine brain. Specifically, this study presents a workflow for whole mouse brain analysis using tissue clearing, LSM and an open-source atlas registration tool for c-Fos mapping to study changes in neuronal activity after diffuse brain injury using the CHIMERA TBI model. We present: 1) SHIELD passive clearing as a method of choice for optimal retention of endogenous c-Fos TdTomato signal, 2) modification of the open-source package, BrainQuant3D, for c-Fos+ cell mapping in the cleared brain image dataset, 3) validation of BrainQuant3D output with an additional method, and 4) multiple visualisation approaches that can be utilised to dissect the large brain-wide region-specific cell count acquisition enabled by whole brain mapping tools.

## MATERIALS AND METHODS

### Animals

All animal procedures were approved by the University of British Columbia Committee on Animal Care (protocol A19-0264) and were performed in compliance with the Canadian Council on Animal Care guidelines. Mice were kept on a reverse 12h light:dark cycle at Centre for Disease Modelling facility with indoor temperature ranging between 20-22°C and humidity between 50-60%. Mice were fed the 5053 diet (PicoLab® Rodent Diet 20; protein diet formulated to limit genetic variability) and had ad libitum access to water and food throughout the entirety of the study. For brain-wide neuronal activity labelling, transgenic reporter mice Fos^2A-iCreER^ (TRAP2) (DeNardo et al., 2019) ; Jackson labs stock # 030323, that express a tamoxifen-inducible Cre recombinase from the *Fos* promoter, were crossed with the Cre-dependent TdTomato reporter mouse line Ai14 (Jackson labs stock # 007914) and maintained as hemizygotes for both transgenes.

### CHIMERA TBI Procedure

Mice (male, 4-5mo old) were anesthetized with isoflurane (induction: 5%, maintenance:3-4%, in oxygen 1L/min) during the entirety of the procedure. Anesthetized animals received lubricating eye ointment to prevent corneal drying, and saline (10mL/kg) and meloxicam (1mg/kg) by subcutaneous injection for hydration and pain control, respectively. Mice were positioned and secured in a supine position on the CHIMERA impactor device (Namjoshi et al., 2014) and anesthesia was maintained through a nose cone until immediately before the head impact. A toe-pinch was performed to confirm a surgical plane of anesthesia before induction of trauma using a stainless-steel piston. Mice were randomised into TBI or sham groups where the TBI group received two mild impacts at 0.5J induced 24 hours apart. Sham mice underwent all procedures except head impact. Animals were placed in a heated recovery cage until consciousness was regained.

### Behavioural assessments

#### Loss of righting reflex

Loss of righting reflex (LRR) latency was measured from the time of impact for TBI animals or cessation of isoflurane for sham animals, until the animal become ambulatory in the recovery cage.

#### Neurological severity score

Acute neurological impairment was assessed using the Neurological severity score (NSS) in all animals within 2 hours of the procedure as described previously (Namjoshi et al., 2014). NSS is a composite of ten tasks that evaluate motor ability, balance, and alertness to assess injury severity and neurobehavioral recovery. One point was awarded for failure on each task up to a maximum NSS of 10. NSS scores classify injury as fatal/near fatal (NSS=9-10), severe (NSS=7-8), moderate (NSS=5-6) or mild (NSS<5).

### Tamoxifen Injection

4-Hydroxytamoxifen (4-OHT; Sigma H6278-10mg) stock solution (20mg/ml) was prepared by dissolving 10mg of 4-OHT in 0.5 ml of 100% ethanol solution by shaking (1300rpm) at 37°C for 20min (Guenthner, Miyamichi, Yang, Heller, & Luo, 2013). In a separate vial, oil mixture was prepared using one part castor oil (S259853, Sigma) to four parts of sunflower oil (S5007, Sigma). 0.5ml of oil mixture was added to the 4-OHT solution and ethanol was evaporated using a vacuum centrifuge until the final volume dropped to 0.5ml. 4-OHT (dose: 50mg/kg) (Guenthner et al., 2013) was administrated intraperitoneally 6 h after injury or sham procedures using a 25g needle. Animals were monitored daily for wellbeing and weight loss in their home cages for 7 days until euthanasia.

### Euthanasia and tissue collection

Mice were anesthetized with intraperitoneal ketamine/xylazine (150/20mg/kg). Upon reaching a surgical plane, blood was collected using cardiac puncture into tubes containing EDTA and animals were then perfused with 20ml of cold heparinized PBS at a flow rate of 8ml/min followed by 30ml of 4% paraformaldehyde (PFA) in PBS. Whole brain tissue was dissected and collected in labelled 20ml glass bottles and post-fixed in 4% PFA for 24hrs at 4°C with shaking.

### Blood biomarker analysis

Whole blood was centrifuged at 1500g for 15 min, plasma was separated and kept at -80°C until assayed. Levels of plasma murine GFAP (glial fibrillary acidic protein) were quantified by a homebrew immunoassay using MesoScale Discovery (MSD) platform as published (Button et al., 2021). Levels of plasma neurofilament light chain (NF-L) were measured by MSD Neurofilament-light assay kit (Mesoscale, cat# K1517XR-2) following the manufacturer’s instructions.

### SHIELD Tissue Clearing

The SHIELD (<u>S</u>tabilization to <u>H</u>arsh conditions via <u>I</u>ntramolecular <u>E</u>poxide <u>L</u>inkages to prevent <u>D</u>egradation) tissue processing and clearing method was used to render mouse brains transparent (Park et al., 2019). Ready-to-use commercial SHIELD preservation and passive clearing reagents were acquired from LifeCanvas technologies (Spin-off company by (Park et al., 2019)). To ensure tissue preservation before delipidation, resins were embedded in PFA-fixed brains by incubating them with 20ml of SHIELD OFF solution (containing SHIELD buffer solution, SHIELD Epoxy solution and distilled water prepared in 1:2:1 ratio respectively) for 4 days at 4°C with shaking. The SHIELD OFF solution was then replaced with SHIELD ON buffer for 24hrs at 37°C with shaking. For clearing, tissues were incubated in 20ml passive clearing buffer at 37°C for ∼3-4 weeks. Samples were visually inspected for the extent of clearing using a light box until they were optically translucent but not transparent. To wash excessive SDS from the clearing buffer, samples were washed with PBST (1% Triton X-100 in 1xPBS) at 37°C overnight with shaking and stored in 1X PBS + 0.02% sodium azide at 4°C until ready for imaging.

### Light sheet microscopy

For optical clearing prior to imaging, the refractive index of the tissue was matched to 1.46 by dipping it in 20ml of 100% EasyIndex solution (LifeCanvas Technologies) overnight at 37°C with gentle shaking. Optically transparent tissue was imaged using a Zeiss Light Sheet Z.1 Microscope System (Z.1 LSM; Carl Zeiss Microscopy, Germany), which is ideally suited for the imaging of thick specimens using separate illumination and detection light paths to minimize photobleaching and maximize imaging depth. For whole brain imaging, the Mesoscale Imaging System (Translucence BioSystems) was used as it adapts the Z.1 LSM to enable imaging of large optically cleared whole-brain tissues with unprecedented speed and resolution, which is not possible with the standard Z.1 chamber. The system consisted of a custom-designed imaging chamber that is 43% larger in volume compared to the Z.1 standard chamber. A 3D-printed specimen holder was used to affix the brain (cerebellum up and olfactory bulb down) to the flat end of the holder to allow immersion of the tissue into the sample chamber that was filled with EasyIndex. A 561 nm laser was used to visualize c-Fos+ cells. The brain was imaged horizontally with a 5X objective and a 0.34 zoom to generate a volumetric 3D stack of the whole brain.

### Image processing

Image datasets acquired using the LSM were processed to generate 3D renderings for downstream analysis. The .czi file from Zen Black software (Zeiss, Germany) was imported into Arivis Vision 4D image analysis software (Zeiss, Germany). Arivis is a modular software that facilitates analysis with multi-channel 2D, 3D and 4D images of >1TB data size and allows stitching and alignment of large multi-dimensional image stacks acquired using LSM. After importing files into Arivis, tiles were stitched together with the image stitcher tool. Tiles were sorted based on acquisition sequence and overlayed using the ∼100 μm overlapping area between two consecutive tiles. Accurate alignment of the tiles was ensured by manually checking every 100^th^ plane in the dataset. Stitched files were then used for 3D rendering in the Arivis 4D module that provides an interactive interface for whole brain volumetric rendering. Movies were generated using the Storyboard feature and exported at 20-30% of the original resolution (given the large size of files). The 2D planer view allowed navigation and .tiff export of the image planes (resembling the traditional imaged sections from the brain) in the plane of interest (i.e., sagittal, coronal or horizontal orientation).

### ClearMap and SMART

ClearMap is a python-based tool designed for analysis of light sheet microscopy data (Renier et al., 2016) from cleared brain and is available at GitHub (https://github.com/ChristophKirst/ClearMap2). SMART is a R based package (Jin et al., 2022) for interactive atlas alignment (https://github.com/sgoldenlab/SMART). Both packages were installed and tested using tutorial data and user instructions.

### BrainQuant3D

BrainQuant3D is a python-based package that allows processing of large-scale microscopy data to segment target cells and generate cell densities by region (https://github.com/sunilgandhilab/brainquant3d). BrainQuant3D was developed for whole-brain imaging analysis of localization patterns of monocytes (Schneider et al., 2019). The BrainQuant3D pipeline utilizes two editable files, *parameter.py* and *process.py*, to specify image features and parameters for the run. We retooled the package by modifying the .py files for estimation of c-Fos+ cells in the mouse brain by optimising a custom ilastik filter. The process of changing filters was not included in the BrainQuant3D tutorial documentation and required examination of the existing code base. A pixel classification filter was made using ilastik 1.3.3 (Berg et al., 2019), and its path was designated in the respective BrainQuant3D file. With the ilastik filter implemented BrainQuant3D segments cells not based on their intensity in the original image, but on the probability ilastik has assigned to a pixel as a measure of how likely that pixel is to be a cell given the classifier trained by ilastik. We used a Bash script to automate processing and tested probability thresholds ranging from 0.1-0.7 to aid in determining a threshold for analysis. After comparing data and visually inspecting the segmentation in Fiji, a probability of 0.30 was used. We also reduced the complexity of the atlas registration transform by using an Affine transform in place of the original SynRNA transform due to holes (i.e., blank areas) appearing in the atlas registration likely due to excessive warping. The modified package has been uploaded to the GitHub (https://github.com/sunilgandhilab/brainquant3d) by forking the original package (https://github.com/MehwishUBC/BrainQuant3D_cFos) and contains imaging and analysis data files presented in this study.

### Life Canvas (LC) mapping and registration

Brains used for method comparison were also prepared using SHIELD passive clearing. Fixed TRAP2 brains were first SHIELD preserved and then were cleared passively at 45°C for 7 days with *Clear+* delipidation buffer followed by Batch labelling with NeuN (Mouse Monoclonal Antibody, Cat# MCA-1B7) using *SmartBatch+* for the atlas registration channel (Life Canvas Technologies). Samples were imaged with SmartSPIM (Light sheet Microscope, LifeCanvas Technologies) using 3.6x objective at 4 μm z-step and 1.8 μm xy pixel size in the transverse plane (Channels: 561 nm – c-FosTdTomato, 642 nm – NeuN). Alignment and cell detection was performed in 3D using SmartAnalytics software (LifeCanvas Technologies). A correlation score was calculated to reflect a good atlas vs. sample alignment that reflects voxel brightness between the warped sample and the atlas. A high value indicates that locations with higher and lower brightness are aligned between the warped sample and the atlas, while a lower value reflects a worse correspondence. For the samples presented here, the correlation score is 782708: 0.75, 782710: 0.82, 781742: 0.81, 787337: 0.81, which is indicative of a good alignment. To provide a visualization of the density per brain region across the brain volume, heat maps were generated using *SmartAnalytics* by Life Canvas Technologies. Each heatmap displays a section of the Allen Brain Atlas, with borders highlighted, colored based on the density value from the corresponding points in region file. The heatmap shows the maximal granularity division of regions as per the Allen Institute Brain common coordinate framework (Allen Institute for Brain Science, 2017). The heatmap plates show every 25 steps in the atlas (625 microns apart) with atlas voxel sizes of 25x25x25 μm. Heatmaps showing averaged absolute c-Fos+ cell density of sham and TBI group as well as subtracted (TBI-sham) averaged c-Fos+ cell density were created.

### Region based 3D analysis

To assess number of c-Fos+ cells within a region of the 3D brain dataset, regions of interest (ROI) were drawn manually in the whole-brain isotropic image dataset (9x9x9um) by using the object drawing tool in Arivis Vision 4D. Hippocampal boundaries were marked by tracing hippocampal anatomical landmarks on 3-5 sections randomly dispersed within the volume of the brain that contains the hippocampus (accuracy confirmed with Allen Brain Institute interactive atlas (https://mouse.brain-map.org/static/atlas). ROI were first drawn in both the sagittal and coronal planes to assess which approach provided better details for anatomical tracing. The coronal plane was thereafter selected for all brains as drawing in the sagittal plane was somewhat compromised due to the separation of hippocampus into two parts on some planes. The Arivis tool used the input ROI and interpolated the complete structure to generate a “volumetric sea horse shaped hippocampus” on each side of the brain. Once rendered, the hippocampus was fed to the analysis pipeline as a 3D object with a custom ROI feature. An object mask was created to eliminate counting of c-Fos+ cells from non-hippocampal regions. The number of c-Fos+ cells within the hippocampus were counted using the Blob finder feature with diameter set to 10 pixels with a probability threshold of 15% and split sensitivity of 30%. An object feature filter was applied to exclude blobs < 4 pixels to eliminate noise. Mapped blobs (c-Fos+ cells) were rendered in 3D and movies with key frames were exported at 35% resolution. The number of c-Fos+ cells in hippocampi from both hemispheres were obtained and mean hippocampal c-Fos+ numbers were plotted for each group (sham or TBI) and correlated with hippocampal region c-Fos+ cells obtained from same brains using BrainQuant3D, Life Canvas and 2D particle analysis.

### Region based 2D analysis

Using ImageJ, the 16-bit 2D images were analyzed by an analyst blinded to the sample groupings. Two consecutive planes were selected at roughly bregma -2.0 mm (dorsal hippocampus) and two consecutive planes were selected at bregma -3.2 mm (ventral hippocampus). ROI were manually drawn as signals were generally between grey intensity 1100 to 2200 and background 450 to 700. c-Fos positivity was defined by the ’Find maxima’ function with prominence set at 500. Signal density was defined as the number of maxima per million pixels. The reported signal density was the average value of the 2 consecutive planes.

### Data visualisation and analysis

Whole brain c-Fos+ cell count data generated via the BrainQuant3D (BQ3D) or the Life Canvas methods were visualised using Arivis (https://www.arivis.com/products/pro), Fiji (https://fiji.sc/; (Schindelin et al., 2012)), GraphPad Prism (https://www.graphpad.com/) and R (https://www.r-project.org/). Sunburst plots were constructed in R using the “sunburstR” and “d3r” libraries by importing the hierarchical c-Fos cell density data from a multilevel structural detail input excel sheet. Interactive sunburst plots were saved as .html files and the code is available with the modified BrainQuant3D package on GitHub ((https://github.com/MehwishUBC/BrainQuant3D_cFos) as well as the Open Science Framework (OSF) page (https://osf.io/2j5bd/). Unsupervised hierarchical clustering analysis on c-Fos+ cell count data from 200 selected brain regions with granularity less than 5 (4 children structure within parent structure) to investigate if the differential c-Fos+ cell counts can cluster cases together. No assumptions were made in clustering analysis, and thus, the “unsupervised” analysis determined relatedness of the cases irrespective of their assigned experimental group. The analysis was carried out in R using the Gplots package.

## RESULTS

### Whole brain clearing and light sheet imaging

To ensure minimal damage to tissue and maximum signal retention, we tested the SHIELD passive clearing method to render the whole murine brain transparent for light sheet imaging (Park et al., 2019). Using TRAP2 mice, which permanently express the immediate early gene c-Fos in neurons upon tamoxifen injection, we optimised buffer quantities, clearing time and incubation temperature (37°C) for an adult (3-4 months old) murine brain with both hemispheres, brainstem and cerebellum intact. The entire process from tissue collection to imaging took 6 weeks. The preservation method allowed excellent c-Fos signal retention without tissue shrinkage or inhomogeneous clearing. The c-Fos+ labelling was distinct in cell bodies as well as in immediate processes (**Movie 1** - showing c-Fos+ labelled ∼1mm thick tile from cleared TRAP2 brain tissue). Furthermore, SHIELD fixation facilitated easy handling of the cleared tissue and mounting to the imaging holder with minimal to no damage or artifacts. We used a custom-made large chamber and sample holders (Translucence Biosystems) that are compatible with the Zeiss light sheet Z1 microscope allowing for imaging of the whole murine brain in one piece. For each brain, the image acquisition time was approximately 4 hours and the acquired image dataset file was approximately 300 GB in size when the brain was imaged at 5X magnification, 1.8 x 1.8 x 4 μm pixel size (xyz), 1250 x 1250 image resolution, using two laser channels (488 nm and 561 nm) with 1250 x1250 image resolution. Increasing the resolution, zoom, or adding channels increased imaging time. Stitching of the acquired volumetric image datasets using image analysis software Arivis allowed a user-friendly interface for 3D rendering of the brain as well image processing for cell segmentation and analysis.

### Mapping neuronal activity after CHIMERA TBI as an exemplar for workflow optimisation

To establish a workflow for generating 3D brain-wide maps using tissue clearing and LSM data, CHIMERA TBI experiments were carried out in TRAP2 mice (n=4). To assess TBI-related alterations in neuronal activity, adult male TRAP2 mice were subjected to two successive CHIMERA TBI injuries 24 hours apart each at 0.5J of kinetic energy in the sagittal plane. 4-OHT was injected 6 hrs after the 2^nd^ TBI to capture acute changes in neuronal activity via Tdtomato tagged c-Fos+ cell distribution **(Figure 1A)**. Acute neurological impairment assessment revealed that, compared to control mice, injured mice took longer to regain their righting reflex indicating injury effects in the TBI group **(Table 1)**. At 7 days post-injury, cardiac blood was collected for biomarker analysis which showed a trend of increased blood GFAP (mean: 17.9 vs 211.45) and NFL levels (mean: 80.75 vs 318.15) in the TBI group **(Table 1)**. Brain tissues were collected and processed for SHIELD, clearing, imaging and 3D rendering **(Movie 2)**. The c-Fos+ cells were distributed throughout the brain including olfactory bulb, hippocampus, cortex, hypothalamus, brainstem and cerebellum **(Figure 1B-D).**

**Figure 1.**
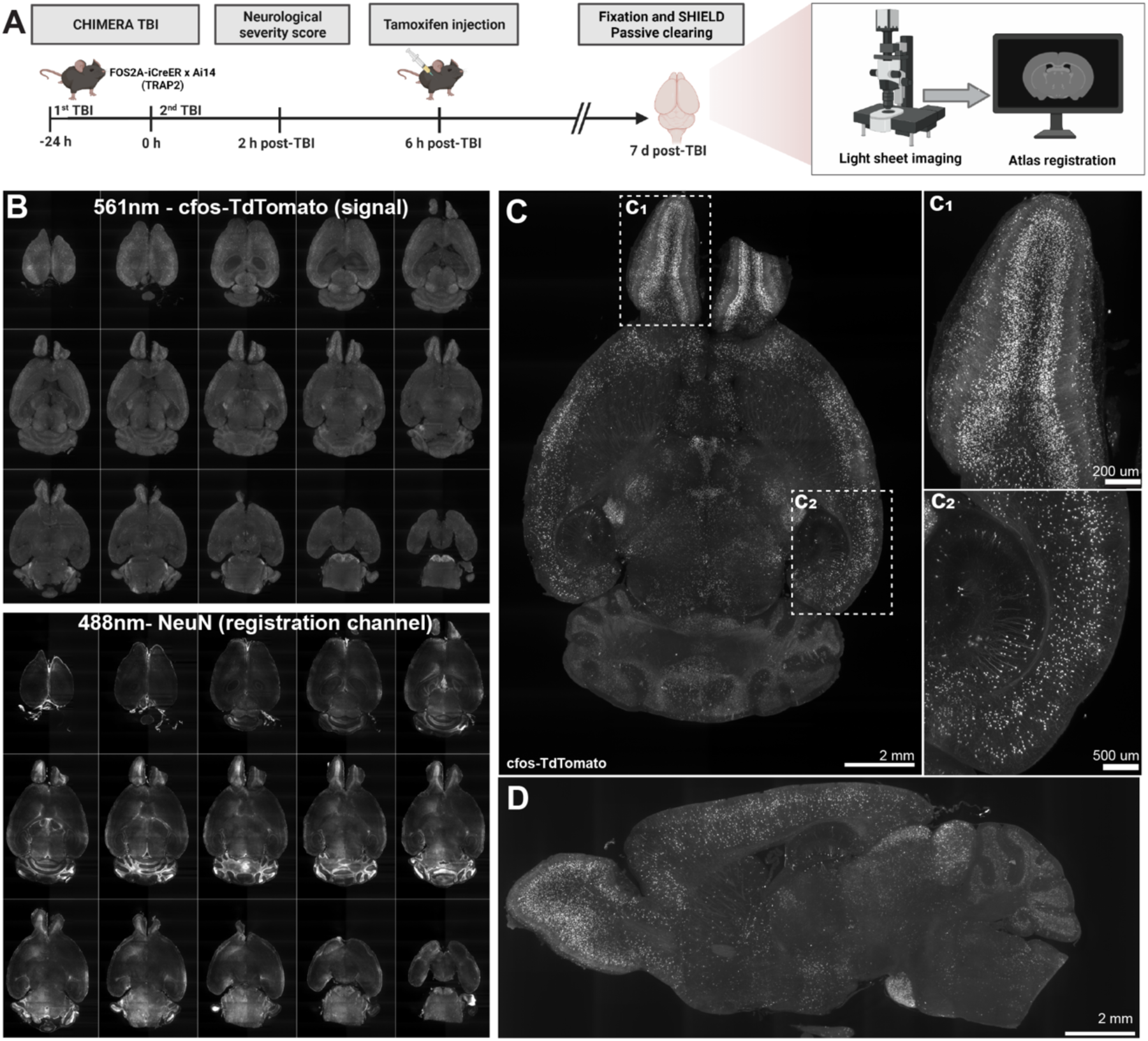
(A) Experimental design. TRAP2 mice were subjected to 2 mild (0.5J) TBI 24 hrs apart or sham procedures in the control group. Neurological severity score was assessed 2 hours after the 2^nd^ TBI and tamoxifen (4-OHT) was injected at 6 hrs after the 2^nd^ TBI to constitutively label c-Fos-expressing cells with TdTomato. At 7 days post TBI, PFA-fixed brains were SHIELD fixed and passively cleared for light sheet imaging, atlas registration, segmentation and regional c-Fos+ cell estimation. (**B-D) Whole-brain montage of a representative sham brain.** Using light sheet microscopy, images were captured from entire intact brains using **(B)** the 561nm laser for c-Fos+ signal and 488nm laser for NeuN labelling (neuronal marker) for atlas registration. **(C-D)** The c-Fos+ cells were distributed throughout the brain including (C_1_) olfactory bulb, (C_2_) hippocampus, cortex, brainstem and cerebellum. (C=horizontal view, D=sagittal view). Scale bars; C=2 mm, C1 200 μm, C2=500 μm and D=2 mm.

**Table 1.** showing age, body weight, duration of exposure to anesthesia before injury as well as duration of loss of righting reflex, neurological severity score and biomarker levels after TBI or sham treatment. Abbreviations: LRR= Loss of righting reflex, NSS=Neurological Severity Score, GFAP= Glial fibrillary acidic protein, NFL= Neurofilament light chain

| Mouse ID | Group | Age (days) | Body weight (g) | Isoflurane duration (sec) |  | LRR duration (sec) |  | NSS (score 1-10, where 10 is most severe outcome) |  |  | GFAP (pg/ml) | NFL (pg/ml) |
| --- | --- | --- | --- | --- | --- | --- | --- | --- | --- | --- | --- | --- |
|  |  |  |  | 1<sup>st</sup> TBI | 2<sup>nd</sup> TBI | 1<sup>st</sup> TBI | 2<sup>nd</sup> TBI | 2 hr after 2<sup>nd</sup> TBI | 1 d after 2<sup>nd</sup> TBI | 2 d after 2<sup>nd</sup> TBI | 7 d after 2<sup>nd</sup> TBI | 7 d after 2<sup>nd</sup> TBI |
| 782708 | Sham | 157 | 42 | 455 | 233 | 181 | 95 | 0 | 0 | 0 | 21.21 | 55.98 |
| 781742 | Sham | 157 | 37 | 372 | 238 | 74 | 84 | 0 | 0 | 0 | 14.74 | 105.50 |
| 787337 | TBI | 143 | 30 | 241 | 250 | 129 | 248 | 0 | 0 | 0 | 325.68 | 444.32 |
| 782710 | TBI | 156 | 41 | 233 | 280 | 84 | 532 | 0 | 1 | 0 | 97.22 | 191.98 |

The image dataset was analysed using two independent atlas registration methods; 1) establishing BrainQuant3D for in-house c-Fos+ cell analysis; and 2) using the Life Canvas contract research organization (CRO) service for c-Fos+ cell analysis to ensure reliability of the open source BrainQuant3D method. To differentiate between the outcomes of the two atlas registration methods tested, data generated by the BrainQuant3D package and the CRO service will be hereafter referred as BQ3D and LC respectively. In the following sections, we first present comparisons between both methods and then describe regional comparisons of c-Fos+ cell counts between sham and TBI groups using LC data as an example of potential of the pipeline to detect brain-wide changes after injury. BQ3D code and its output files are available on GitHub and Open Science Framework (OSF) page with guiding notes to enable users with minimal coding skills to carry out analyses.

### Brainquant3 was retooled for brain-wide c-Fos+ cell segmentation and atlas registration

Compared to traditional 2D histological cell counting approaches, 3D whole brain imaging datasets require automated and robust tools to segment labelled cells throughout the brain while identifying the specific location within the multi-millimeter tissue image set. Therefore, after optimising the clearing and imaging of the intact murine brain, we next tested open-source packages for atlas registration and cell segmentation.

Commercial imaging analysis software typically embed image processing and analysis pipeline including compression, reconstruction, and segmentation of datasets. Some examples of popular commercial software include Arivis Vision4D (which is also utilized in this workflow for 3D rendering), Amira, Imaris, and Image-Pro Plus. However, none of these tools so far offer warping and registration with a mouse brain atlas which eventually limit their utility for an entire workflow. Moreover, the price of these software packages motivated us to find an open-source tool that allows warping and atlas registration thereby enabling whole brain analysis similar to in-vivo MRI imaging pipelines. However, the challenge with open-source tools is the lack of maintenance from developers as well as need for fluency in coding in various languages.

We tested ClearMap (Renier et al., 2016), SMART (Jin et al., 2022) and BrainQuant3D (Schneider et al., 2019) for our analysis with little success using the first two. SMART (Semi-Manual Alignment to Reference Templates) is an R package that builds a pipeline which interfaces with the *Wholebrain* package (Fürth et al., 2018) and offers a console interface that allows image adjustment and manual correction for registration errors. However, despite resampling our data to match the dimensions, physical scaling, and intensity scaling of the tutorial dataset, SMART was unable to consistently segment brain from background using the regi_loop function. During the autoloop stage of regi_loop, the *Wholebrain* package is used for a first pass of atlas plate registration on the experimental data, and successful completion of the autoloop is required for subsequent manual correction. If regi_loop were to run successfully, we planned to quantify the number of planes that needed manual correction alongside the required degree of correction as a key performance metric for SMART. However, the autoloop stage repeatedly failed and was unable to fully process our dataset, making performance quantification and cell segmentation not possible.

BrainQuant3D is a python-based package (Schneider et al., 2019) that was developed as a modification of ClearMap (Renier et al., 2016) to handle large datasets obtained using the Z1 LSM (Zeiss, Germany). While we were able to successfully run BrainQuant3D using the tutorial data for monocyte segmentation, the code required modifications to enable segmentation of c-Fos+ cells in our imaging datasets **(Figure 2A)**.

**Figure 2.**
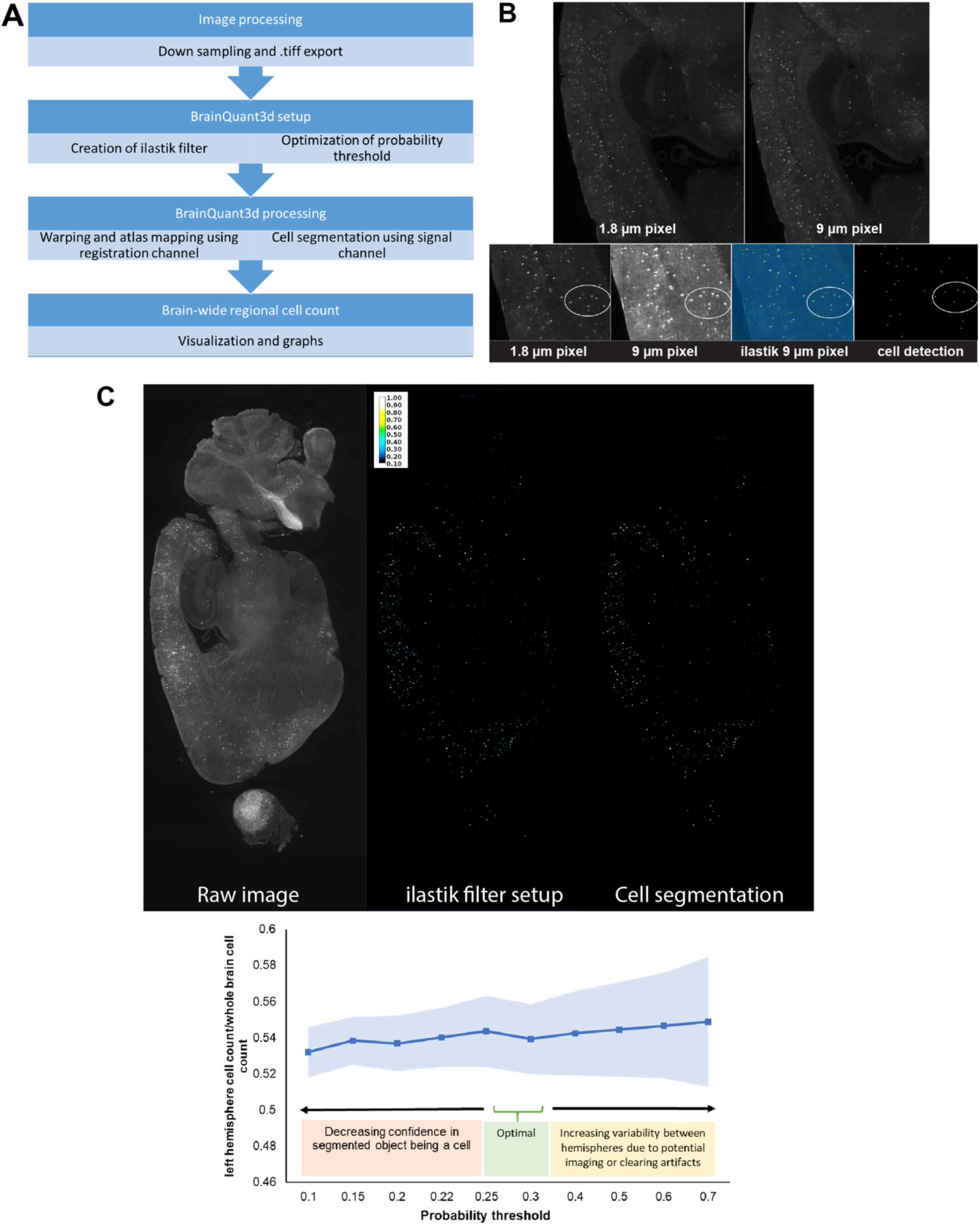
BrainQuant3D for brain-wide c-Fos+ cell segmentation. (A) BrainQuant3D workflow. Imaging data acquired using LSM is first processed by downsampling and export of the stitched images as .tiff files. Next, the BrainQuant3D filter is established by creating a custom ilastik filter that segments c-Fos+ cells and probability threshold optimisation to reliably measure cell counts throughout the brain. Within the BrainQuant3D processing step, the registration channel image set is used for warping and atlas mapping whereas cell segmentation is carried out using the c-Fos signal channel. Finally, regional c-Fos+ cell counts are acquired as a .csv file and then analysed via data visualisation tools. **(B) Downsampling.** Changing resolution of images to 9 x 9 x 9 μm pixel size did not result in loss of information and was efficient to establish the ilastik filter and c-Fos+ cell segmentation. **(C) Probability thresholding.** The probability of 0.30, i.e., pixels of an intensity with 0.30 or higher are considered as cells, showed the best cell segmentation whereas a probability lower or higher than 0.30 resulted in inaccurate estimation of c-Fos+ cell counts in brain due to artifacts or noise. Note: Image brightness is enhanced for illustration purpose only.

We performed the analysis one hemisphere at a time in the sagittal plane using the respective Allen Institute Mouse brain atlas. Firstly, we downsampled images to match their pixel dimensions of the tutorial data i.e., 9 x 9 x 9μm in xyz planes **(Figure 2A-B)**. Importantly, downsampling did not result in loss of information or distortion of images **(Figure 2B)**. Secondly, we found that the rolling background subtraction filter provided with the package was ineffective for c-Fos+ cell segmentation. Consequently, we created a custom ilastik filter created using our dataset **(Figure 2C)**. The process of changing filters was not included in the BrainQuant3D tutorial documentation and required examination of the existing code base. Therefore, a pixel classification filter was made using ilastik 1.3.3 (Berg et al., 2019), and its path was designated in the respective BrainQuant3D file. Lastly, we also reduced the complexity of the atlas registration transform by using an Affine transform in place of the original SynRNA transform due to holes (i.e., blank areas) appearing in the atlas registration image output that was likely due to excessive warping. A Bash script was written for basic automation of processing, and a probability of 0.30 was used **(Figure 2C)** after comparing results between the probabilities as well as with outcomes acquired using another analysis method that is further discussed below.

### Brain-wide c-Fos expression using BrainQuant3D and Life Canvas registration methods

To confirm the output of BrainQuant3D method, we acquired atlas registration and c-Fos segmentation of TRAP2 sham and TBI brains on the same image dataset using the Life Canvas (LC) CRO service using their *SmartAnalytics* tools. The whole brain c-Fos+ cell counts with all regions included acquired by both methods showed a linear relationship for each brain with R^2^ values ranging between 0.86-0.97 **(Figure 3A).** Regional comparison also identified comparable trend in c-Fos+ cell counts for each brain **(Figure 3B)**.

**Figure 3.**
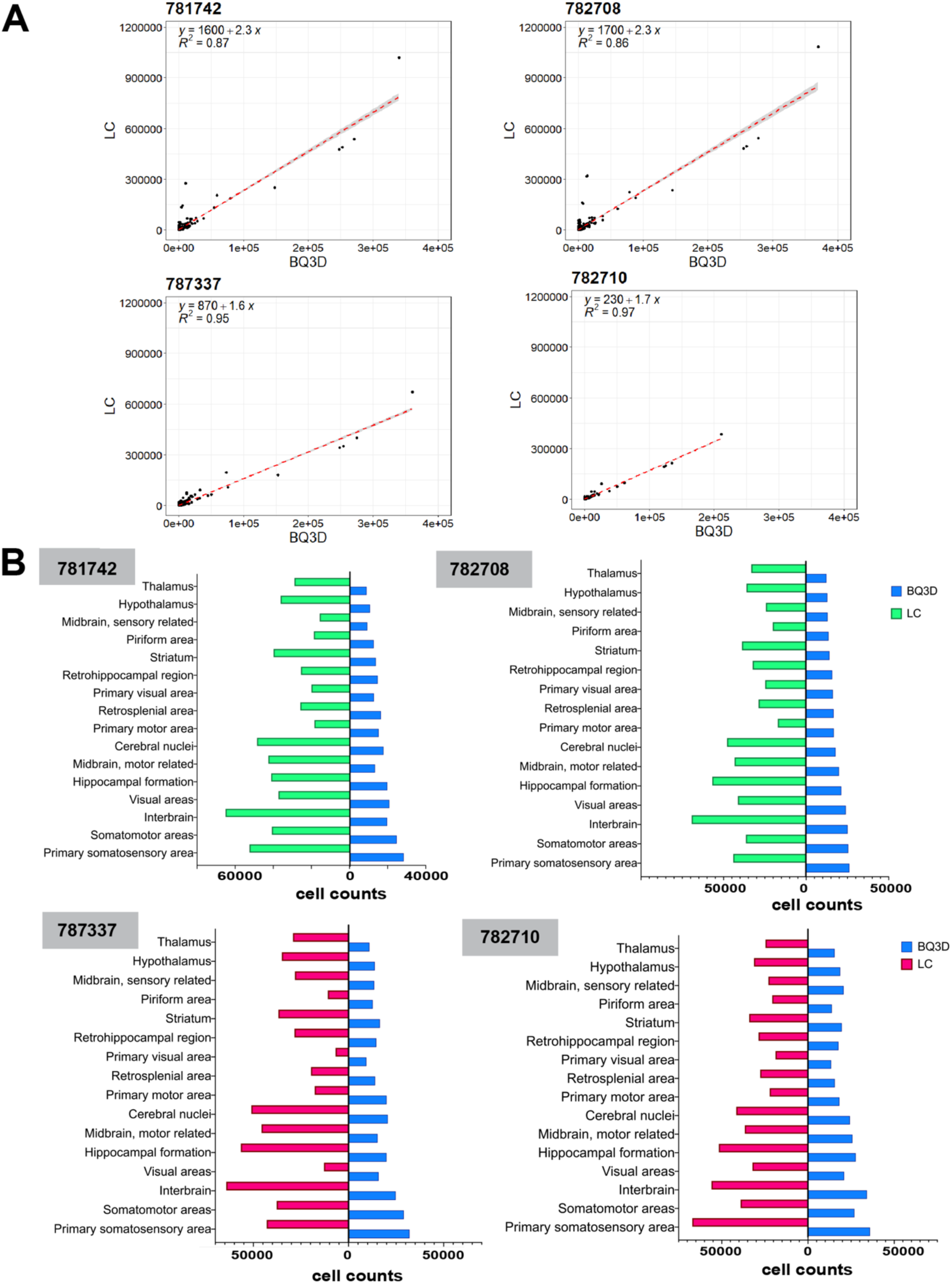
BrainQuant3D (BQ3D) vs Life Canvas (LC) c-Fos+ counts. **(A)** Correlation plots showing c-Fos+ cell counts from each brain using two independent methods. **(B)** Horizontal bar charts showing regional c-Fos+ cell counts for each brain acquired by BQ3D and LC. Numbers in grey boxes indicate study ID of each mouse brain.

### Region based analysis for method validations

As both BrainQuant3D and Life Canvas registration methods are automated and lack a user interface for verification of mapping accuracy, we next validated the c-Fos+ cell counts in the hippocampus through two ROI based methods carried out by two independent analysts:

#### a) Region based 2D particle analysis

Dorsal and ventral hippocampal ROIs were manually drawn on exported .tiff images by a rater blinded to study ID and group assignment **(Figure 4 A)**. The total c-Fos+ cell counts from both dorsal and ventral hippocampal areas were compared for all brains. Compared to sham controls, the TBI group showed a decrease in c-Fos+ cell density consistent with observation from the BrainQuant3D and Life Canvas quantification methods that utilised automated scripts to estimate these numbers. The coefficient of variance (CoV) estimated for 2 consecutive planes in the same region at the same laterality was 15%. The number of c-Fos+ cells estimated using 2D particle analysis corelated with the number of c-Fos+ cells acquired from BQ3D and LC **(Fig 4C)**.

**Figure 4.**
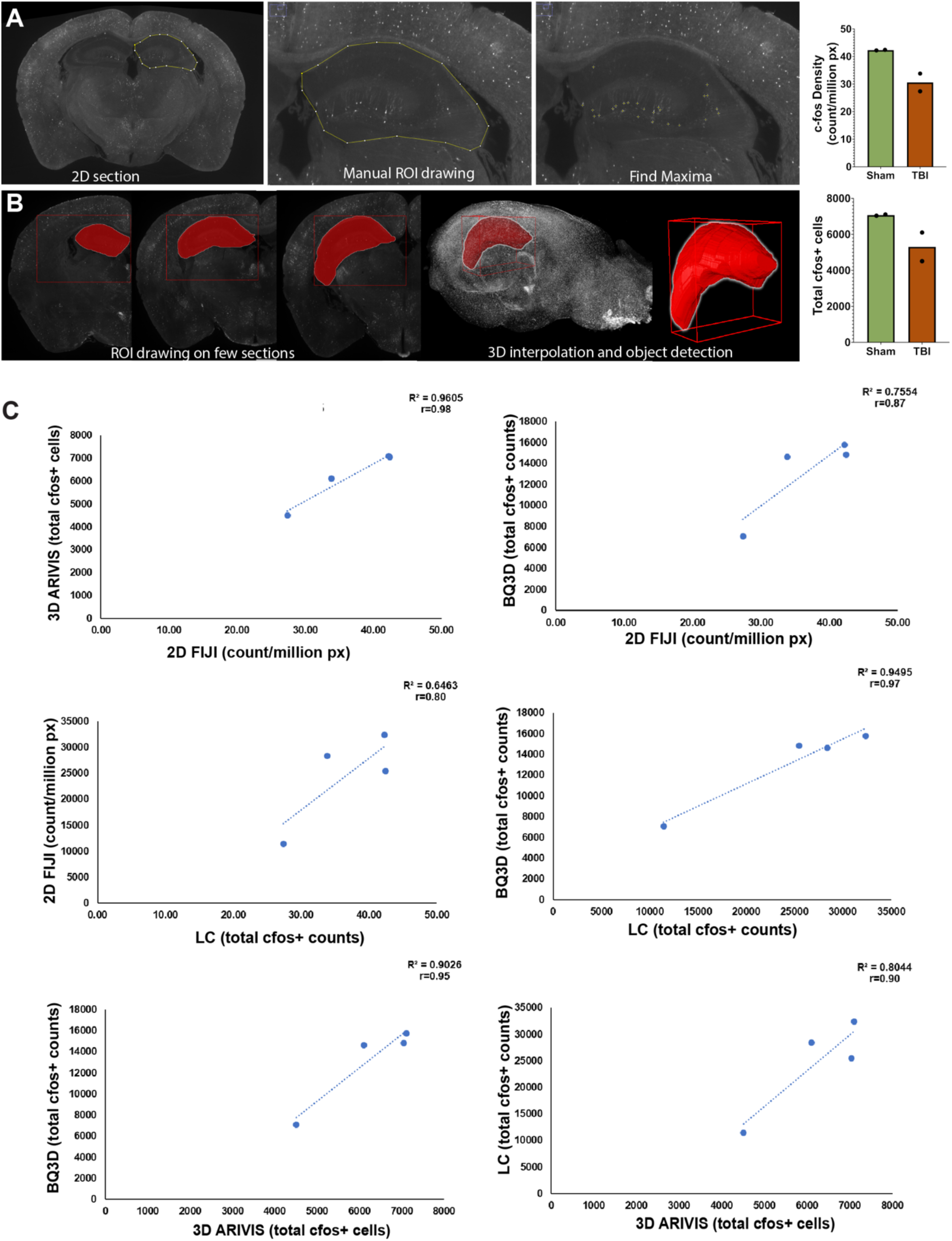
Regional analysis for method comparison. **(A) 2D particle analysis.** ROI was manually drawn around the dorsal and ventral hippocampus on traditional 2D images of sections (representing planes) in the Fiji tool. The number of c-Fos+ cells/million pixels were determined using the “find maxima” feature in Fiji (yellow cross). **(B) 3D particle analysis**. ROI encompassing whole hippocampus was constructed by marking anatomical boundaries of dorsal and ventral hippocampal segments on 3-5 sections randomly dispersed along the anterior-posterior aspect of the brain that were then interpolated by Arivis to form the complete hippocampus on each side of the brain. The “blob finder” feature of Arivis counted all c-Fos+ cells (blobs) in the hippocampus. **(C) Comparison between methods.** The c-Fos+ cell estimates in hippocampus were comparable between methods utilised for estimation. Scatter plots showing c-Fos+ counts/brain for all four brains with R^2^ and Pearson correlation coefficient (r) value on each plot.

#### b) Region based 3D particle analysis

To validate the number of c-Fos+ cells acquired using BrainQuant3D, a 3D interactive ROI based approach in Arivis was also used to account for volumetric estimates **(Figure 4 B, Movie 3)**. The number of c-Fos+ cells estimated using the Arivis Blob finder corelated with the number of c-Fos+ cells acquired from BQ3D and LC **(Fig 4C)**. In addition to validation of the BrainQuant3D counts, this approach can also be used for a specific regional analysis given that the anatomical landmarks are easily delineable.

### 3D brain-wide analysis as a unique tool to identity spatial neuronal activity signatures after TBI

Acquisition of whole brain spatially enriched neuronal activity data allowed a detailed investigation of changes caused by TBI. Spatial heatmaps were generated to present cell density across the brain volume using annotated Allen Brain Atlas sections (625 μm apart) **(Figure 5)**. Overall, c-Fos+ cell density was lesser within the TBI group compared to sham controls (3015.5 ± 759 vs 4685.2 ± 205) **(Figure 5A)**. Mean subtracted c-Fos+ cell density of the groups highlighted multiple brain areas with reduced c-Fos+ cells including somatosensory cortical areas, hippocampi, several brain stem and cerebellar cortical areas.

**Figure 5.**
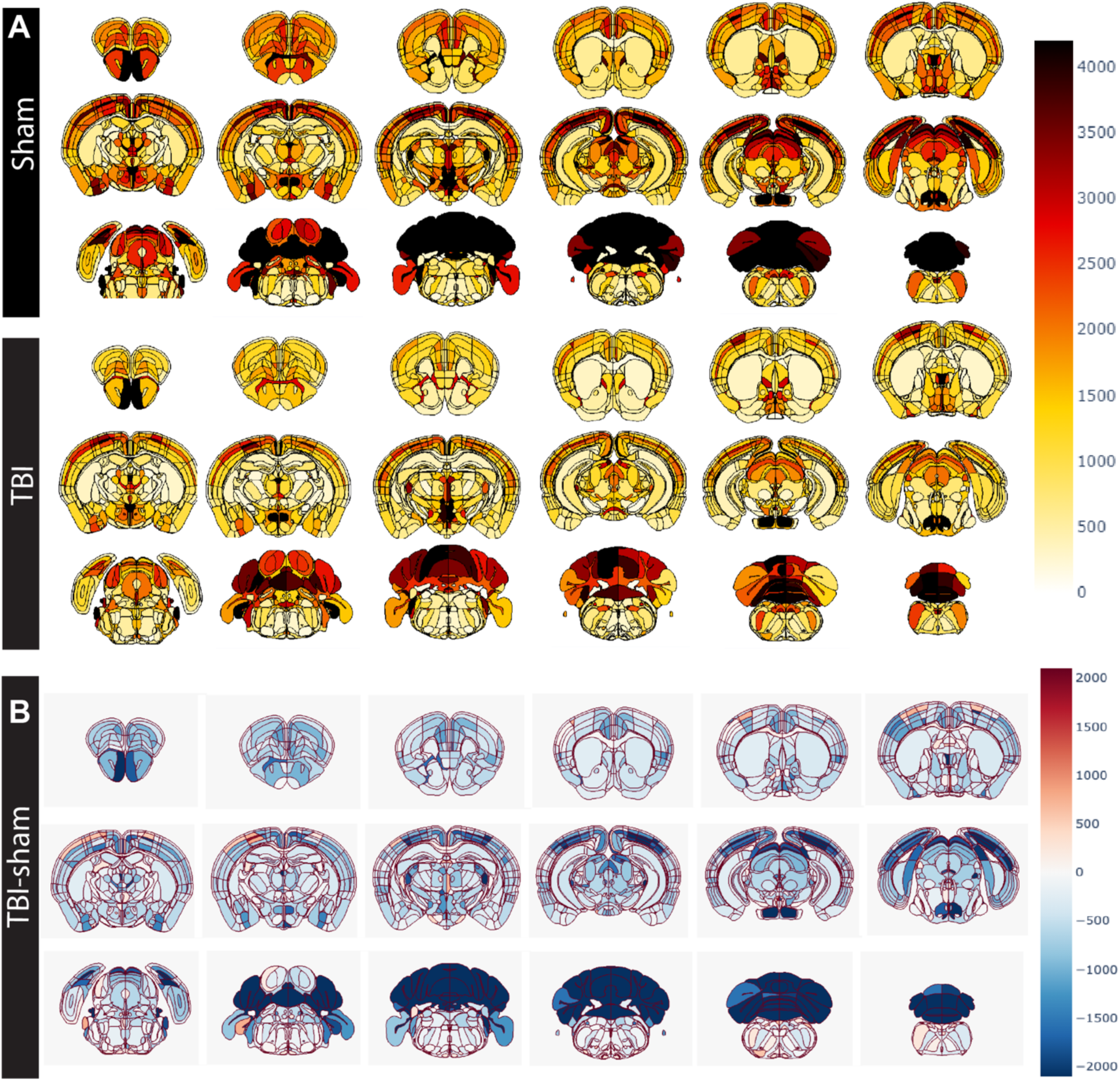
Spatial distribution of c-Fos+ cells across brain regions. **(A)** Heatmaps showing mean c-Fos+ cell density on Allen Mouse Brain Atlas planes (625 microns apart) for sham (upper panel) and TBI (lower panel) groups. **(B)** Heatmaps showing subtracted (TBI-sham) mean c-Fos+ cell density. Note brain-wide reduction in c-Fos+ cell density across brain regions in the TBI group with marked decreases in cerebral and cerebellar cortices.

### Brain-wide multi-level data visualisation allows comparative analysis

To facilitate in-depth data visualisation and comparative analysis, we also established a code in R to plot hierarchical regional c-Fos+ cell density data following Allen Mouse Brain Atlas annotation. These plots are interactive and can be used as a visual interface for regional comparisons (**S1-2**). In our case study, mean c-Fos+ cell density in the cerebral cortex of the TBI group was 2,996 ± 822 compared to 4,443 ± 35 in the cerebral cortex of the sham group. In hippocampus, the mean c-Fos+ cell density in the dentate gyrus of the TBI group is 1630 ± 717.89 compared to 951 ± 422.2 in the sham group **(Figure 6, S1-2)**. Hippocampal region c-Fos+ cell density in dentate gyrus constituted 98.6% of the total hippocampal c-Fos+ cell density in sham group and 93.7% of total hippocampal c-Fos+ cell density in TBI group **(Figure 6)**. From the ∼1600 regional cell counts/brain dataset, regions with granularity up to 5 (parent structure with 4 children structures) were extracted and log2Fold change between sham and TBI groups were estimated **(Figure 7, S4)**. Overall, an increase in cfos expression was observed in hypothalamic regions whereas thalamic and hippocampal regions exhibited reduced cfos expression after TBI **(Figure 7)**. We also carried out an unsupervised hierarchical clustering analysis to determine if the regional c-Fos+ cell count can segregate the cases into distinct clusters. No assumptions were made in clustering analysis, hence the unsupervised analysis determined the relatedness between cases without information regarding their experimental group assignment. As a result, the clustering analysis identified two main clusters in the data where cluster 1 grouped both sham cases together and cluster 2 contained both TBI groups, indicating robustness of c-Fos+ cell count in identifying sham from TBI **(S3)**.

**Figure 6.**
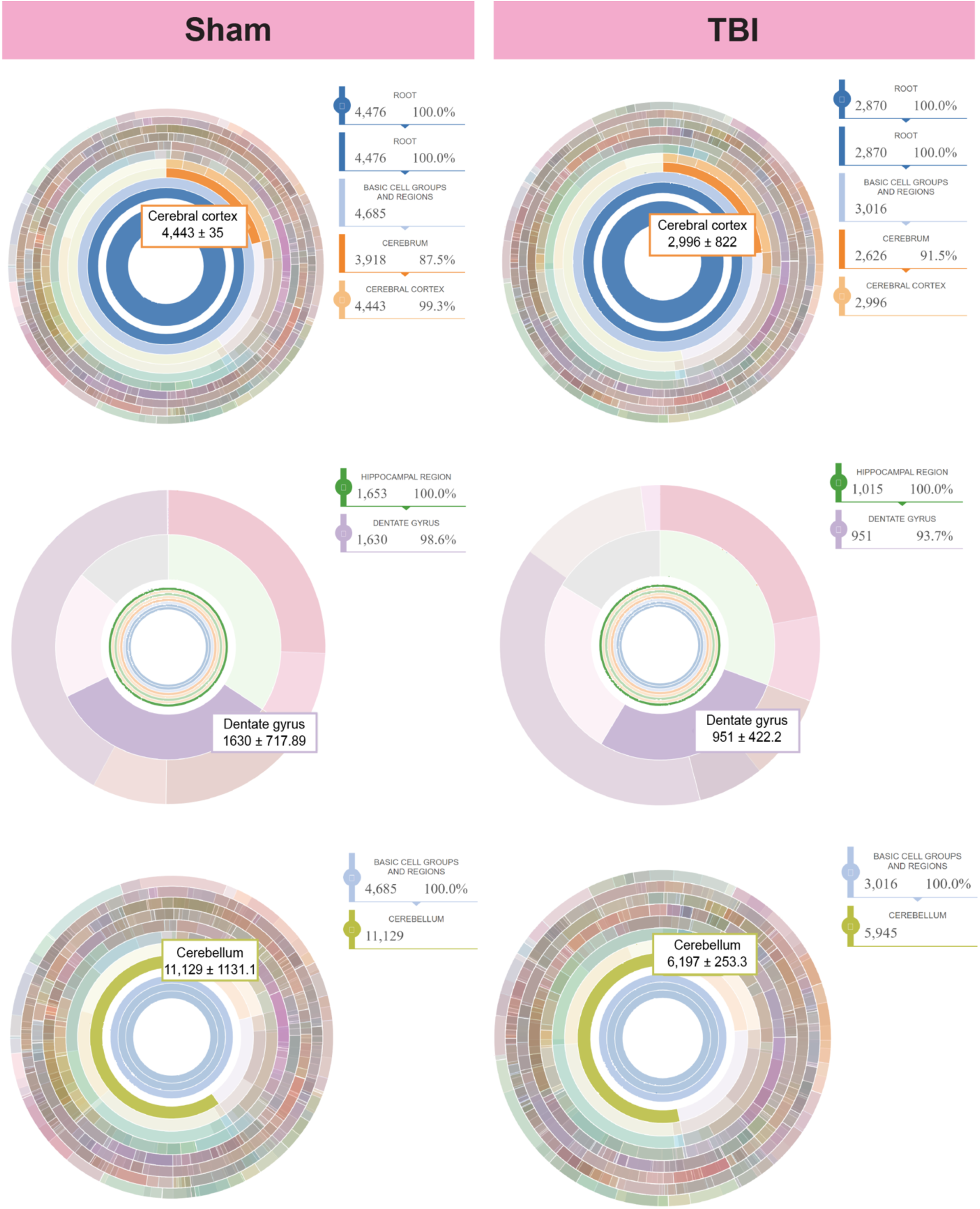
Brain-wide multi-level data visualization. Sunburst plots created using R showing mean regional cell density i.e., number of c-Fos+ cells/regional volume (cells/mm^3^) as rings of the “sunburst” with its hierarchical structural relationship in the averaged sham and TBI whole brain datasets. Inserts on each sunburst plot present the mean cell density (± standard deviation) of the region within the experimental group (sham on left and TBI on right) and sequential legends on the side of each sunburst plot show percent contribution of c-Fos+ cell density within the region level. In the sunburst plot, the innermost ring represents whole brain cell counts and the subsequent ring represents *basic cell groups and regions* (as annotated by the Allen Mouse Brain Atlas) with consecutive rings representing cerebrum, brainstem and cerebellum regions i.e., each subsequent ring represents child structures of the parent structure. All regions follow the annotation and hierarchical structure of the Allen Mouse Brain Atlas and “arc length” is proportional to percent cell density of region within the larger ring. Plots are interactive and hovering over a ring shows regional identity, hierarchical level, and percent cell density of region within the ring. Interactive .html files for each group are included in supplementary materials, GitHub page and OSF project.

**Figure 7.**
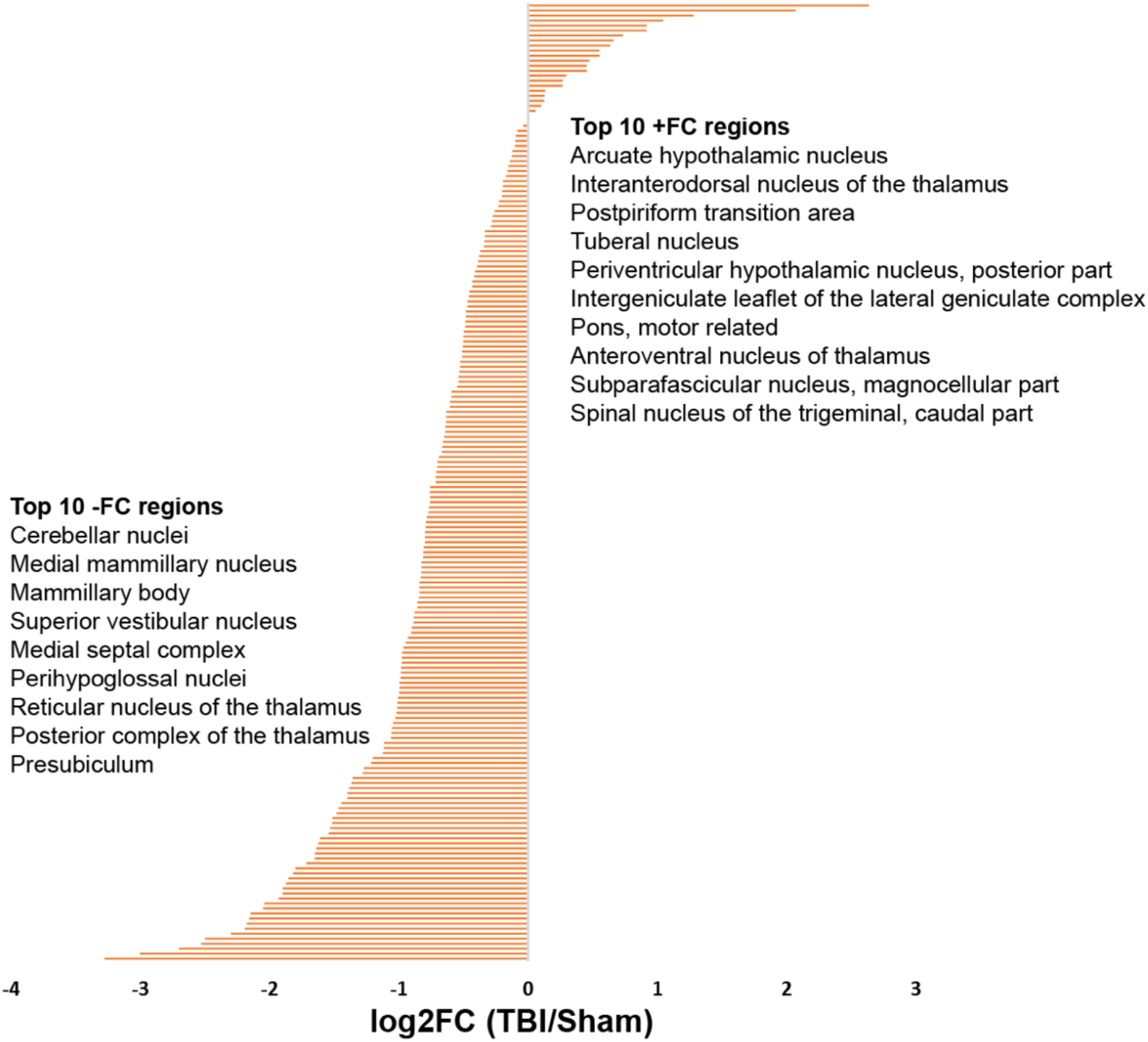
Differential c-Fos+ cell count between sham and TBI. From the whole-brain data, regions with granularity level up to 5 (4 children structure within parent structure) were selected and log2Foldchange was calculated (TBI/sham). Brain regions were sorted by highest (+) to lowest (-) Log2FC values and plotted as a horizontal bar chart. The top 10 regions with highest +FC and -FC are shown. The complete list is provided in Supplementary Materials (S4).

### Whole brain analysis provides opportunity for studying network-wide changes in brain

Next, we demonstrate the utility of this 3D whole brain mapping workflow as an unbiased pan-regional analysis tool to identify network wide changes. Here, we show the effect of TBI on brain-wide c-Fos expression as an example of dysregulated neuronal activity and potentially functional connectivity. We utilised the homologous functional network annotation from resting-state functional magnetic resonance imaging (rs-fMRI) comparative data on functional connectivity in rodent and human brain (Xu et al., 2022). We identified a TBI-induced differential regulation in c-Fos+ cell density within the default mode network, limbic network, lateral cortical network, motor and auditory network, visual network, and somatosensory network **(Figure 8)**.

**Figure 8.**
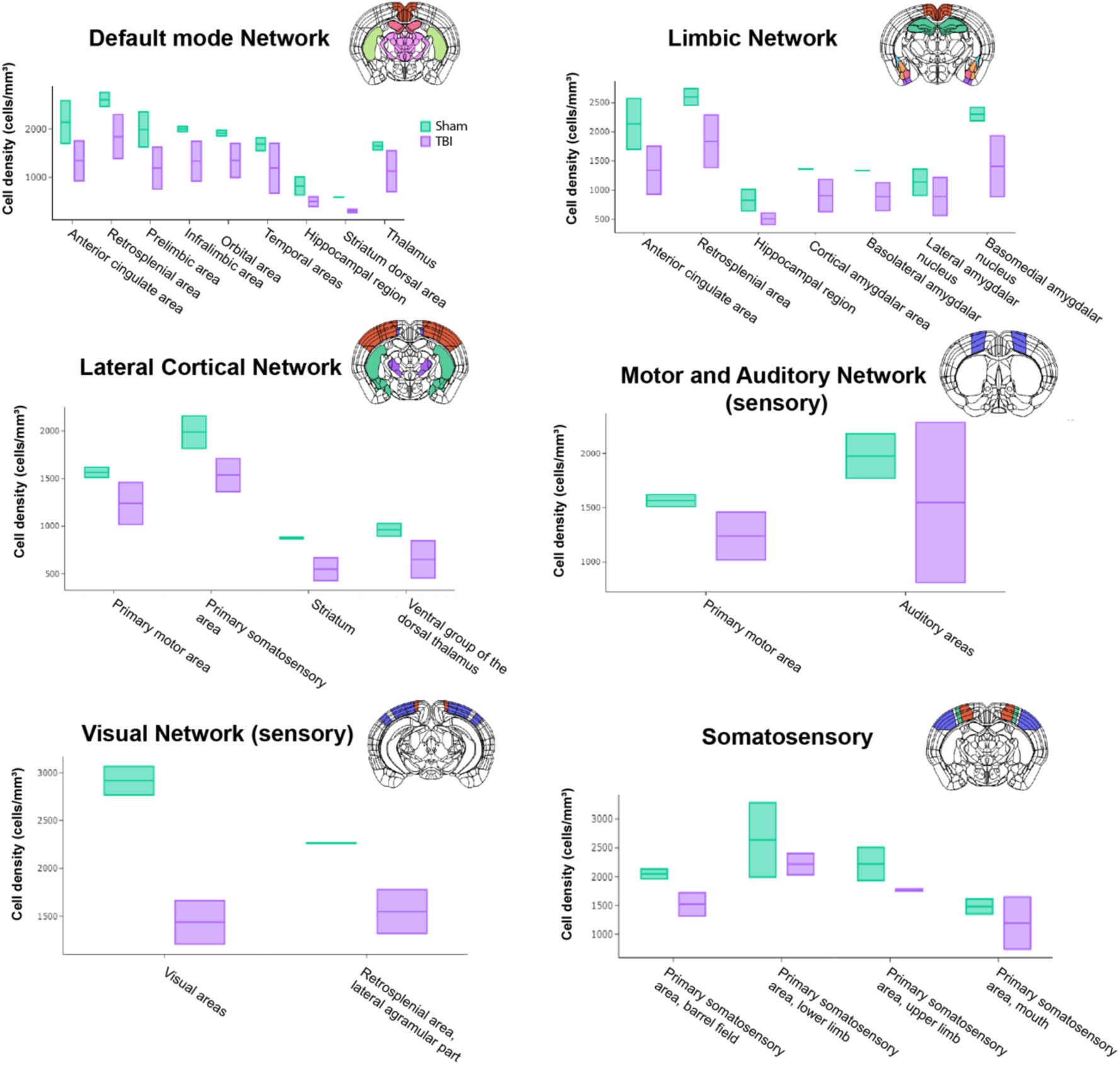
Brain-wide functional connectivity. Whole brain analysis provides an opportunity for studying network-wide changes after TBI. Graphs illustrate the ability of 3D brain neuronal activity mapping to identify changes in the default mode, limbic, lateral cortical, motor and auditory, visual, and somatosensory networks in the murine brain after TBI. Sub-regional analysis can predict functional implications of network changes. Data are presented as c-Fos+ cell density from sham and TBI (n=2/group) brains, where the upper and lower bar represents individual density and the middle bar representing the average of both brains. Representative atlas icons depict anatomical location of regions within the network.

## DISCUSSION

In this study, we optimised a workflow for whole murine brain clearing, imaging and cell segmentation with atlas registration and demonstrate that this 3D segmentation analysis approach serves as a versatile tool for mapping brain-wide spatiotemporal changes after TBI. The methods described here are applicable to other neurological disease models in mice.

The SHIELD clearing approach used here resulted in robust signal retention of TdTomato labelled c-Fos cells and required 4-6 weeks of incubation in the clearing. Clearing time depends both on the size of the tissue and the incubation temperature, which can be increased to 45°C to accelerate clearing albeit this may result in reduced compatibility with immunolabelling. More recent approaches suggest the potential to reduce clearing time through use of an electrophoretic apparatus (Hee Yun et al., 2019). Eventually, the choice of clearing method depends on the robustness of the signal, size of tissue, imaging system compatibility and extent of immunolabelling (Richardson & Lichtman, 2015). We used a modified imaging chamber (Translucence Biosystems) to enable imaging the of the intact murine brain. However, the need to immerse tissue in the refractive index matching solution limited the availability of compatible immersion-based higher magnification optics. More recently available microscopes such as MesoSPIM (Voigt et al., 2019) may enable greater flexibility in imaging with higher resolution.

BrainQuant3D, a version of ClearMap, was developed for immune cell segmentation in mouse brain (Schneider et al., 2019). To segment c-Fos+ cells effectively, we modified the package and designed a custom ilastik filter optimised using our imaging dataset as well as identified the efficient probability thresholding parameters for representative cell segmentation. These modifications enable the use of this atlas registration and mapping tool to be adaptable for a wide variety of cell-like brain wide analysis after imaging. Recently, the FASTMAP (Flexible Atlas Segmentation Tool for Multi-Area Processing) image analysis tool was developed as an open-source tool platform to facilitate atlas registration and densitometry analyses (Terstege, Oboh, & Epp, 2022). However, FASTMAP requires selection of 2D mouse brain atlas planes to identify most-closely matched image of immunolabelled sections followed by ROI drawing and manual free form resizing of ROI corresponding to the atlas. While this method accounts for shrinkage of tissue from histological processing, it requires considerable time to manually draw regions and requires the user to properly identify neuroanatomical regions. By contrast, BrainQuant3D provides segmented image output files as well as whole hemisphere counts within ∼30 minutes, which is a considerable advantage given the wealth of unbiased data generated with this method.

In this workflow, we used the Arivis Vision4D image analysis software due to its highly user-friendly non-coding interface for 3D rendering of the whole brain image dataset. This can be partially acquired using open-source tools such as Fiji. Furthermore, the BrainQuant3D cell segmentation and atlas registration package that we optimised in this study is not limited by the upstream processing tool used as long as the input files are tiffs with 9x9x9 μm pixel size. The package description states that the orientation of the input files can be changed as needed (e.g. sagittal, coronal or horizontal orientation) with a compatible atlas annotation file. However, we found that the sagittal orientation was most efficient in the warping and segmentation processes that can be also attributed to the image resolution in each plane.

Overall, there are a wide range of methods available for cleared tissue analysis depending on experimental needs. Manual ROI-based cell counting is not ideal when seeking region-specific whole brain cell counts due to laborious and highly time-consuming processes. By contrast, while some commercial software or CROs can streamline the analysis process, offer flexibility, and provide powerful tools, their cost makes them inaccessible for many researchers. Consequently, open-source, fully automated pipelines such as BrainQuant3D possess benefits of both time-efficiency and cost-effectiveness, making them highly attractive for brain-wide analysis.

The findings presented here are from 4 whole brain assessments. While each brain contained 800 rows of data/hemisphere (totalling to 6400 values of regional c-Fos+ cell counts), differences observed between sham and TBI groups need further validation. The data generated in this pipeline testing study will guide future power calculations as well as raise new hypotheses for how TBI affects neuronal activity and connectivity. Using a similar workflow with iDISCO+ clearing and light sheet imaging, Frankowski et al 2022., presented a brain-wide map of input to inhibitory neurons in a mouse model of TBI (Frankowski et al., 2022). However, the atlas registration method they used requires manual annotation of each cell position in Cell Finder and a multi-step process for atlas registration using *Brainreg* and *ImageJ* tools. While feasible for assessment of cell counts in magnitudes of a few hundred, these methods will be virtually impossible for estimation of brain-wide counts that range in hundreds of thousands. BrainQuant3D provides the unique advantage of acquiring cell counts from about 800 parent and child regions (annotated by Allen Mouse Brain atlas) per hemisphere in about 20 minutes.

Taken together, the data presented here allow decrease potential sampling bias and facilitate the identification of diffuse pathological changes in brain after TBI, changes in functional networks. We demonstrate that diffuse mild TBI leads to differential dysregulation of multiple functional networks as shown by decrease in c-Fos+ cell density, a neuronal activity marker, in regions involved within these networks. For instance, the default mode network (DMN) encompasses functional connectivity with cortical areas including cingulate, retrosplenial, prelimbic, infralimbic and orbital areas as well as hippocampal and thalamic regions. The DMN “connectopathy” has been shown in animal models of Autism Spectrum Disorder (ASD) (Pagani et al., 2021) and Alzheimer’s disease (AD) (Adhikari, Belloy, Van der Linden, Keliris, & Verhoye, 2021) similar to aberrant functional connectivity in human attention-deficit/hyperactivity disorder, ASD and AD populations (Lim et al., 2014; Uddin et al., 2008; Uddin, Yeo, & Spreng, 2019). These insights help in hypothesis generation as well as identification of novel pathological implications of diffuse brain injury that can validated by studies designed with in-depth network analysis. Recent advances in *ex vivo* imaging modalities and computational pipelines integrating whole brain imaging approaches with *in vivo* imaging emphasize wide applicability of the workflow presented here (G. A. Johnson et al., 2023). In conclusions, we present here a feasible workflow with the power to map brain-wide changes after experimental TBI in mice and that is applicable to other neurological disease investigations using murine models.

## ACKNOWLEDGMENTS

The authors would like to thank Anna Wilkinson and Parsa Alizadeh for their overall support to the project. The authors also thank Hesham Soliman (UBC), Nicole Cheung (UBC), Courtney Akitake (Zeiss), Damian Wheeler (Translucence Biosystems), and Brian Nguyen (LifeCanvas) for discussions on imaging setup and data visualisation. This work was supported by resources made available through the Dynamic Brain Circuits cluster and the NeuroImaging and NeuroComputation Centre at the UBC Djavad Mowafaghian Centre for Brain Health (RRID:SCR_019086).

## AUTHORS’ CONTRIBUTIONS

MA was responsible for study conception, designing experiments, performing all experiments, imaging, analyzing and interpreting results, and preparing the manuscript. MA, JL, SW and AW carried out the computational setup and analysis with supervision from TM and CLW. MA, HC, CB and JF contributed to CHIMERA experiments, tissue collection and processing. WHC performed image analysis and contributed to data analysis. MC provided transgenic mice and expert knowledge in study design and execution. MB assisted in 3D visualisation of data. FR provided access, training and support for light sheet imaging. CLW, MC, PAC, and TM are the supervisory authors on the project. CLW was involved in the concept formation and manuscript preparation. The author(s) read and approved the final manuscript.

## DATA AVAILABILITY

All the data presented in the study is available open-access via GitHub (https://github.com/MehwishUBC/BrainQuant3D_cFos) and Open Science Framework (OSF) channels (https://osf.io/2j5bd/). Imaging data set (.tiff files) from all four brains as well as the ilastik filter are also made available to facilitate adoption of the modified forked package.

## MOVIES

**Movie 1**: Light sheet imaging showing c-Fos+ cells in a ∼2mm think area of the TRAP2 cleared brain.

**Movie 2:** 3D rendering showing c-Fos+ cells throughout the cleared TRAP2 brain in multiple orientations.

**Movie 3:** 3D rendering showing c-Fos+ cell segmentation in hippocampal 3D ROI in whole brain imaging dataset from TRAP2 mice

## SUPPLEMENTARY MATERIALS

**S1 and S2**: Sunburst plot .html files

**S3:** Hierarchical clustering analysis plot

**S4:** Fold change value from 200 regions

