## Supplementary figures and images for "LEVERAGING THE POWER OF 3D BRAIN-WIDE IMAGING AND MAPPING TOOLS FOR BRAIN INJURY RESEARCH IN MURINE MODELS"

### S3_hclust count sup.tif

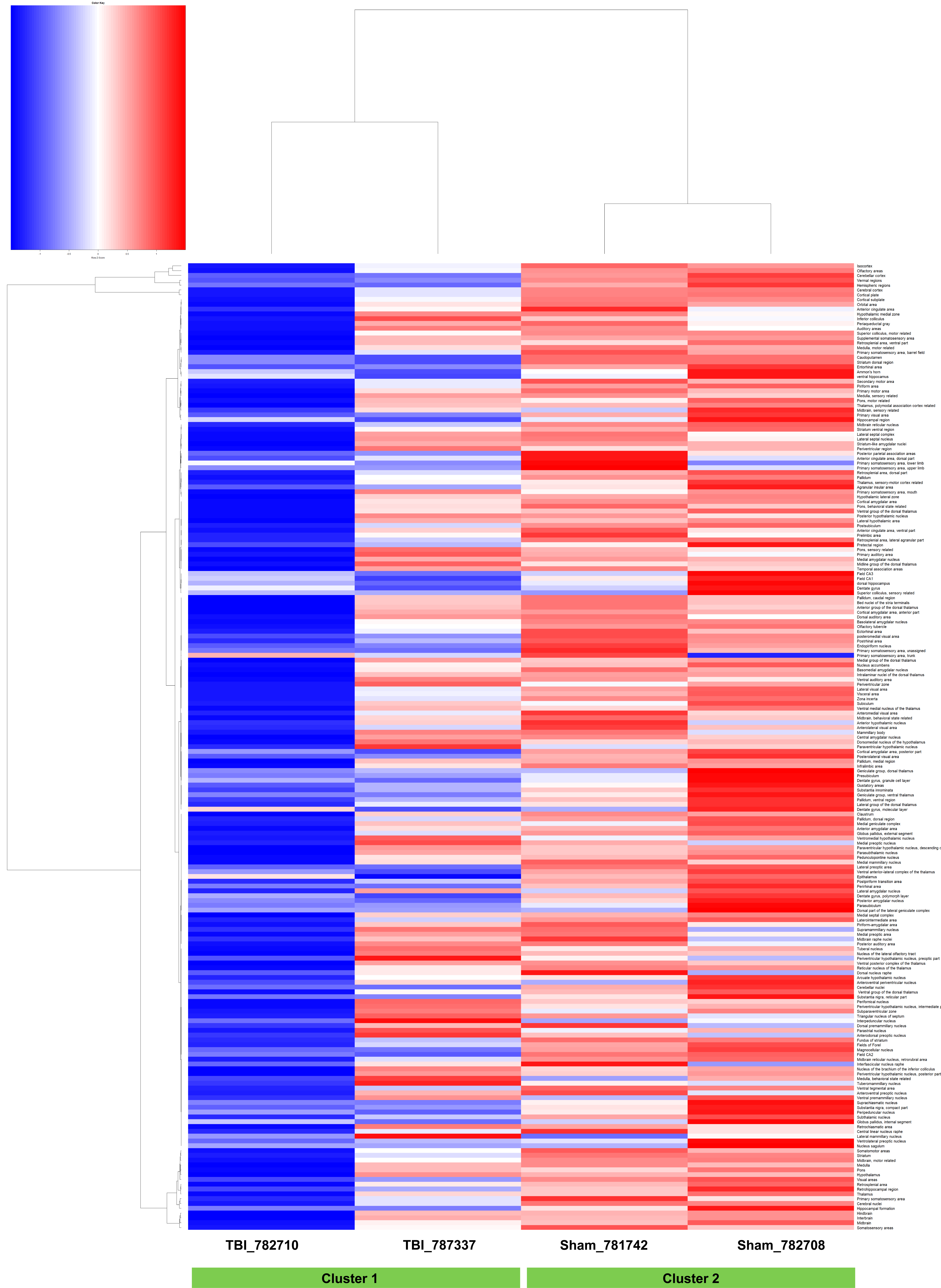
