## Supplementary File 4 for "LEVERAGING THE POWER OF 3D BRAIN-WIDE IMAGING AND MAPPING TOOLS FOR BRAIN INJURY RESEARCH IN MURINE MODELS": S4_FC list.pdf

| Region name | log2FC |
|---|---|
| Arcuate hypothalamic nucleus | -3.27798 |
| Interanterodorsal nucleus of the thalamus | -3.00432 |
| Postpiriform transition area | -2.70079 |
| Tuberal nucleus | -2.52875 |
| Periventricular hypothalamic nucleus, posterior part | -2.49923 |
| Intergeniculate leaflet of the lateral geniculate complex | -2.30051 |
| Pons, motor related | -2.19193 |
| Anteroventral nucleus of thalamus | -2.17681 |
| Subparafascicular nucleus, magnocellular part | -2.15952 |
| Spinal nucleus of the trigeminal, caudal part | -2.1472 |
| Dorsomedial nucleus of the hypothalamus | -2.05048 |
| Medullary reticular nucleus, dorsal part | -2.04029 |
| Anteromedial nucleus | -1.92969 |
| Cerebellar cortex | -1.90072 |
| Medial amygdalar nucleus | -1.89912 |
| Subparafascicular nucleus, parvicellular part | -1.87337 |
| Lateral visual area | -1.85506 |
| Cerebellum | -1.81849 |
| Dorsal part of the lateral geniculate complex | -1.80058 |
| Field CA2 | -1.71672 |
| Posterior hypothalamic nucleus | -1.6521 |
| Cortical amygdalar area | -1.65038 |
| Primary visual area | -1.6405 |
| Paraventricular nucleus of the thalamus | -1.62194 |
| Visual areas | -1.60923 |
| Gracile nucleus | -1.54344 |
| Ectorhinal area | -1.53118 |
| Posterior auditory area | -1.51734 |
| Pons | -1.51425 |
| Paraventricular hypothalamic nucleus | -1.48261 |
| Supratrigeminal nucleus | -1.4668 |
| Medulla, sensory related | -1.44501 |
| Anterolateral visual area | -1.39992 |
| Supraoptic nucleus | -1.39855 |
| Central lateral nucleus of the thalamus | -1.38483 |
| Nucleus of the brachium of the inferior colliculus | -1.36883 |
| Periventricular zone | -1.3592 |
| Parabrachial nucleus | -1.28079 |
| Basomedial amygdalar nucleus | -1.26988 |
| posteromedial visual area | -1.21479 |
| Field CA3 | -1.20201 |
| Hindbrain | -1.12535 |
| Gustatory areas | -1.11776 |
| Entorhinal area | -1.11474 |
| Pontine central gray | -1.06092 |
| Lateral amygdalar nucleus | -1.05795 |

| | |
|---|---|
| Basolateral amygdalar nucleus | -1.05419 |
| Posterior amygdalar nucleus | -1.04601 |
| Endopiriform nucleus, dorsal part | -1.02826 |
| Piriform area | -1.01954 |
| Prelimbic area | -1.01424 |
| Hypothalamic lateral zone | -1.01177 |
| Posterolateral visual area | -1.0078 |
| Pretectal region | -0.99873 |
| Posterior auditory area, layer 5 | -0.99548 |
| Anteromedial visual area | -0.99482 |
| Midline group of the dorsal thalamus | -0.9885 |
| Anterior pretectal nucleus | -0.98534 |
| Orbital area, lateral part | -0.98399 |
| Globus pallidus, external segment | -0.97701 |
| Medial geniculate complex | -0.97497 |
| Central amygdalar nucleus | -0.97311 |
| Red nucleus | -0.96459 |
| Piriform-amygdalar area | -0.94738 |
| Postsubiculum | -0.92861 |
| Precommissural nucleus | -0.90531 |
| Pons, behavioral state related | -0.89964 |
| Ventral auditory area | -0.88643 |
| Infralimbic area | -0.88436 |
| Dorsal premammillary nucleus | -0.87875 |
| Thalamus, sensory-motor cortex related | -0.85776 |
| Primary somatosensory area, nose | -0.85758 |
| Cerebral cortex | -0.84638 |
| Cortical plate | -0.84132 |
| Anterior cingulate area | -0.84131 |
| Subparaventricular zone | -0.8406 |
| Pontine reticular nucleus | -0.8312 |
| Agranular insular area | -0.82766 |
| Retrosplenial area | -0.8265 |
| Endopiriform nucleus | -0.8258 |
| Cerebrum | -0.81298 |
| Medulla | -0.81048 |
| Intermediodorsal nucleus of the thalamus | -0.80703 |
| Midbrain reticular nucleus | -0.79944 |
| Pontine reticular nucleus, caudal part | -0.79847 |
| Orbital area | -0.79766 |
| Ventral tegmental area | -0.79277 |
| Isocortex | -0.79175 |
| Periventricular region | -0.78308 |
| Parvicellular reticular nucleus | -0.77243 |
| Orbital area, ventrolateral part | -0.76658 |
| inferior cerebellar peduncle | -0.76049 |
| Caudoputamen | -0.75966 |

| | |
|---|---|
| Striatum dorsal region | -0.75966 |
| Superior colliculus, optic layer | -0.75844 |
| Primary auditory area | -0.71502 |
| Auditory areas | -0.71491 |
| Laterodorsal tegmental nucleus | -0.71223 |
| Nucleus of the posterior commissure | -0.70473 |
| Anterodorsal nucleus | -0.7021 |
| Lateral hypothalamic area | -0.69174 |
| Thalamus | -0.66932 |
| Central medial nucleus of the thalamus | -0.66534 |
| Paragigantocellular reticular nucleus | -0.66074 |
| Brain stem | -0.65053 |
| Striatum | -0.64104 |
| Secondary motor area | -0.63967 |
| Interbrain | -0.63386 |
| Visceral area | -0.63363 |
| Pons, sensory related | -0.63309 |
| Thalamus, polymodal association cortex related | -0.60555 |
| Hypothalamus | -0.60408 |
| Anteroventral preoptic nucleus | -0.59442 |
| Superior colliculus, sensory related | -0.59301 |
| Ventromedial hypothalamic nucleus | -0.5502 |
| Superior colliculus, motor related | -0.53865 |
| Retrohippocampal region | -0.53784 |
| Somatomotor areas | -0.53219 |
| Primary somatosensory area, lower limb | -0.52587 |
| Mediodorsal nucleus of thalamus | -0.52582 |
| Cerebral nuclei | -0.50999 |
| Midbrain, motor related | -0.50989 |
| Ammon's horn | -0.50587 |
| Orbital area, medial part | -0.50258 |
| Striatum ventral region | -0.50212 |
| Superior colliculus, superficial gray layer | -0.49738 |
| Ventral anterior-lateral complex of the thalamus | -0.48656 |
| Ventral group of the dorsal thalamus | -0.48567 |
| Primary somatosensory area, barrel field | -0.48216 |
| Supplemental somatosensory area | -0.48091 |
| Primary somatosensory area | -0.46939 |
| Nucleus accumbens | -0.46929 |
| Ventral posterior complex of the thalamus | -0.45969 |
| Somatosensory areas | -0.45538 |
| Primary somatosensory area, upper limb | -0.43368 |
| Ventral posteromedial nucleus of the thalamus | -0.4314 |
| Primary motor area | -0.41584 |
| Lateral preoptic area | -0.41171 |
| Anteroventral periventricular nucleus | -0.39376 |
| Lateral septal nucleus | -0.38736 |

| | |
|---|---|
| Midbrain reticular nucleus, retrorubral area | -0.37929 |
| Primary somatosensory area, mouth | -0.37011 |
| Hippocampal formation | -0.34232 |
| Medulla, motor related | -0.33957 |
| Intermediate reticular nucleus | -0.33567 |
| Midbrain | -0.33521 |
| Medial preoptic area | -0.28628 |
| Ventral posterolateral nucleus of the thalamus | -0.27784 |
| Dorsal auditory area | -0.27268 |
| Anterior hypothalamic nucleus | -0.25772 |
| Periventricular hypothalamic nucleus, preoptic part | -0.22996 |
| Spinal vestibular nucleus | -0.22512 |
| Substantia nigra, reticular part | -0.20217 |
| Frontal pole, cerebral cortex | -0.20086 |
| Taenia tecta | -0.19499 |
| Lateral septal complex | -0.19477 |
| stria terminalis | -0.17137 |
| Midbrain, behavioral state related | -0.15961 |
| Hypothalamic medial zone | -0.15376 |
| Periventricular hypothalamic nucleus, intermediate part | -0.14235 |
| Hippocampal region | -0.126 |
| Ventral medial nucleus of the thalamus | -0.12242 |
| Dorsal peduncular area | -0.10042 |
| Midbrain, sensory related | -0.09996 |
| Primary somatosensory area, trunk | -0.09659 |
| Medial group of the dorsal thalamus | -0.08312 |
| Pallidum | -0.03954 |
| Subiculum | -0.00544 |
| Field CA1 | -0.00105 |
| Lateral group of the dorsal thalamus | 0.056079 |
| Lateral posterior nucleus of the thalamus | 0.10007 |
| Inferior colliculus | 0.123398 |
| Magnocellular nucleus | 0.125531 |
| Supramammillary nucleus | 0.130494 |
| Lateral dorsal nucleus of thalamus | 0.26586 |
| lateral forebrain bundle system | 0.26588 |
| Anterior amygdalar area | 0.293452 |
| Medial vestibular nucleus | 0.451469 |
| Parasubiculum | 0.452187 |
| Motor nucleus of trigeminal | 0.472345 |
| Presubiculum | 0.550566 |
| Dentate gyrus | 0.551705 |
| Epithalamus | 0.632177 |
| Posterior complex of the thalamus | 0.658756 |
| Reticular nucleus of the thalamus | 0.731116 |
| Perihypoglossal nuclei | 0.914458 |
| Medial septal complex | 0.916346 |

| | |
|---|---|
| Superior vestibular nucleus | 1.043186 |
| Mammillary body | 1.276614 |
| Medial mammillary nucleus | 2.067453 |
| Cerebellar nuclei | 2.635575 |
